## Supplementary Materials for "LabOS: The AI-XR Co-Scientist That Sees and Works With Humans"

**Supplementary Methods and Notes.**

**S1. LabSuperVision (LSV) benchmark, VLM evaluation prompt, parameter, and raw data.**

For the data collection, each session primarily focused on egocentric AR/XR captures using Smart Glass (e.g. Viture, UReality, or Xiaomi), and DJI Osmo Action Pro 5 cameras. The dataset is partitioned by scene to evaluate generalization across laboratory environments, and includes a balanced distribution of correct and error-containing sessions to support both protocol understanding and failure detection benchmarks. Trained domain experts created and validated annotations. The dataset is split by scene and operator to test generalization and contains equal proportions of videos with and without errors.

The table below describes the key summary statistics for the LabSuperVision (LSV) benchmark data collection process.

***Table S1.1*: LabSuperVision dataset statistics by scene, operator, duration, and issue taxonomy for error detection, correction.**

| Capture equipment | Laboratory Scenes | Video counts | Operator counts | Protocol counts | Issue counts (for error detection, correction) |
|---|---|---|---|---|---|
| **Smart glasses** | Bench (Biomed and Material) | 35 | 8 | 12 | 11 |
| | Functional room (e.g. tissue culture, clean room) | 80 | 8 | 16 | 8 |
| **Action cameras** | Bench | 49 | 7 | 7 | 9 |
| | Functional room (e.g. tissue culture) | 78 | 7 | 11 | 8 |
| **Total** | | **242** | **15** | **46** | **36** |

**The VLM evaluation Prompt details are described below:**

Protocol Generation:
"""
"""
Instructions:
**1. Reference-based direct scoring**
- Compare the “Proposed protocol” against the “Ground truth.”
- Assign a single integer score from **0 (completely unfaithful) to 5 (perfectly faithful)**.

**2. Rubric**
5: All essential steps and details are present, accurate, and in the correct order.
4: Major steps are covered; only minor details or wording differ.
3: Workflow is generally captured but some key steps or details are missing or slightly incorrect.
2: Significant omissions or errors in multiple steps, affecting overall clarity.
1: Most content is missing, misrepresented, or lacks clarity.
0: NOT related to the biology at all. ONLY give this score for empty response or sorry response.

**3. Bias mitigation & reproducibility**
- **Do not** provide any commentary—only output the score.

Output format:
- **Only** return the integer score (0–5), with no additional text.

Ground truth:
{ground_truth}

Proposed protocol:
{proposed}

Score (0–5):
"""
"""

Error detection:
"""
Instructions:
**1) Reference-based direct scoring**
- Compare the “Predicted errors” against the “Ground-truth errors.”
- Assign a single integer score from 0 (completely incorrect) to 5 (perfect).

**2) Matching rules (use these when deciding correctness)**
- Error type: treat close synonyms as a match (e.g., "didn't change tips" ≈ "no tip change").
- Timing: accept approximate matches; a missing/None time in either list is OK if the event clearly corresponds.
- One-to-one is preferred, but one predicted error may cover a single gold error even if phrased differently.
- Hallucinations (predicted errors not supported by ground truth) must reduce the score.

**3) Rubric (0–5)**
5: All gold errors are identified correctly (types align semantically); no hallucinated errors.
4: Most gold errors are correctly identified; at most one minor miss or one minor hallucination.
3: Partial match: some gold errors found but clear misses and/or multiple minor hallucinations.
2: Few gold errors found or several incorrect/hallucinated errors; overall unreliable.
1: Predicted set mostly wrong or dominated by hallucinations.
0: Not related to the task (e.g., empty/sorry/irrelevant) OR gold has errors but none are identified.

4) Special case: when the ground truth has NO errors
- If the prediction also has NO errors → score 5.
- If the prediction hallucinates any errors → score 0–2 depending on severity (use 0 for many or major hallucinations).

5) Bias mitigation & reproducibility
- Do NOT include any commentary—only output the integer score.
- Do NOT infer extra context beyond the provided lists.

Output format:
- Only return a single integer score (0–5), with no additional text.

Ground-truth errors (JSON list):
{ground_truth_errors}

Predicted errors (JSON list):
{predicted_errors}

Score (0–5):
"""

The table below shows the raw data statistics from the evaluation, corresponding to the main text Figure 3.

***Table S1.2*: LabSuperVision Benchmark Enables Evaluation of Leading VLMs for Multi-modal Intelligence in Laboratory Research.**

| **Model** | Protocol Matching Score - GPT-5 eval (Scores 0-5↑) | Error Detection, Correction - GPT-5 eval (Scores 0-5↑) | Protocol Matching Score - human expert eval (Scores 0-5↑) | Error Detection, Correction - human expert eval (Scores 0-5↑) |
|---|---|---|---|---|
| Gemini 2.5 Pro | 1.60 (0.15) | 1.75 (1.29) | 2.40 (0.06) | 1.97 (0.12) |
| Cosmos-Reason-1 | 0.60 (0.12) | 0.81 (0.91) | 2.20 (0.15) | 0.63 (0.09) |
| QWen 2.5-VL 7B | 0.50 (0.13) | 1.06 (0.85) | 1.93 (0.12) | 1.17 (0.12) |

| | | | | |
|---|---|---|---|---|
| GPT-4o | 1.24 (0.09) | 1.06 (1.06) | 1.83 (0.17) | 2.03 (0.03) |

- **Metric: GPT-5 evaluate (left) and human expert evaluation (right). Mean(SEM) shown in bracket.**
- **Prompt and scoring details in the Supp Section (with score rubrics 0-5)**
- **Models.** Gemini 2.5-Pro, Cosmos-Reason-1, QWen 2.5-VL, GPT-4o.

### S2. LabOS Vision-Language-Model (VLM) fine-tuning for lab intelligence

For Fine-tuning a Vision–Language Model for Lab Operations, we implemented the following methods for the VLM within the LabOS physical lab module.

#### Base model and overall training stages

LabOS VLM is a family of multimodal vision–language models (VLMs) adapted for laboratory environments, built upon the Qwen-VL architecture. We trained multiple scales of LabOS VLM (7B, 32B, 72B, and 235B parameters) with both visual and textual encoders jointly optimized for scientific scene understanding. All models were initialized from publicly released Qwen-VL checkpoints and fine-tuned using Low-Rank Adaptation (LoRA) for parameter-efficient adaptation.

#### Datasets

Training utilized two primary multimodal corpora:

1. **FineBio** — a curated physical-lab video dataset containing 38,200 video clips annotated with expert-verified protocol steps, reagent identities, and error labels.
2. **JoVE (Journal of Visualized Experiments)** — 12,000 peer-reviewed video protocols across molecular biology, cell culture, and chemical synthesis domains, converted into frame-level image–text pairs with structured action labels.

Each video was segmented into 10-second clips with synchronized textual descriptions derived from the original narration and annotation schema. The combined dataset was split into 80/10/10 for training, validation, and held-out in-house testing, respectively. The evaluation set included unseen experimental setups to assess generalization to novel laboratory contexts.

#### Supervised Fine-Tuning (SFT)

We first applied supervised fine-tuning (SFT) using LoRA to efficiently update 1–2% of model parameters. SFT aligned multimodal representations on paired experimental video frames and textual explanations covering protocol generation, error detection, and equipment identification. The loss function combined cross-entropy on text generation and image–text contrastive alignment.

Training was conducted with batch size = 128, learning rate = 2e-5, cosine decay schedule, and LoRA rank = 16. SFT training ran for 3 epochs on 8 H100 80GB GPUs using mixed precision.

#### Reinforcement Learning via Group Relative Policy Optimization (GRPO)

To further align model reasoning with expert laboratory preferences, we employed Group Relative Policy Optimization (GRPO) following the framework of DeepSeek-R1. GRPO replaces the explicit critic network of standard PPO with group-relative baseline estimation, reducing computational cost and stabilizing optimization.

For each question q, GRPO samples a group of outputs {o1, o2, ..., oG} from the old policy and optimizes the policy model by maximizing:

J_GRPO = E_q [ (1/G) * sum_i ( min( (pi_theta / pi_old) * A_i, clip((pi_theta / pi_old), 1-epsilon, 1+epsilon) * A_i ) - beta * D_KL(pi_theta || pi_ref) ) ] where A_i = (r_i - mean(r_group)) / std(r_group), and r_i denotes the rule-based reward. We used epsilon = 0.2, beta = 0.05, and G = 8.

#### Reward Modeling

Following DeepSeek-R1-Zero, LabOS VLM adopts a rule-based reward system tailored for experimental reasoning:

- **Accuracy Reward:** evaluates whether the model's generated instruction matches the ground-truth procedural step or correction label (e.g., detecting an incorrect pipetting sequence).
- **Format Reward:** enforces structured reasoning, requiring the model to output its rationale between and tags before emitting the final action.

Rewards are normalized to zero mean and unit variance within each batch. No neural reward model was used to avoid reward hacking and reduce training cost.

#### Evaluation

Model performance was assessed on a held-out in-house dataset containing 2,000 annotated experimental video clips spanning pipetting, transfection, centrifugation, and labeling procedures. Metrics include:

- **Protocol Quality Score (0–5):** human-rated coherence and correctness of generated protocols.
- **Error Detection Accuracy (%):** proportion of video segments correctly classified as compliant or violating the intended protocol.

### S3. LabOS XR-AI Real-time Integration System for Laboratory Operations

#### 1. System Architecture Overview

The LabOS system comprises three interconnected parts operating in concert:

##### 1.1 Capture and Transport (On-device + Edge)

An extended reality (XR) glass (spanning different formats, such as augmented reality/AR, virtual reality/VR, mixed reality/MR), optionally with six-degree-of-freedom (6-DoF) tracking, eye tracking, hand gesture capabilities. The XR headset/glass supports live camera streams, with synchronized video, audio, and optionally with inertial measurement unit (IMU) and related sensors. The smart glass/device system supports user interaction via edge computing or cloud computing infrastructure, through a containerized pipeline utilizing Real-Time Streaming Protocol (RTSP) and gRPC protocols. Capture feeds could optimally include fixed cameras and sensors in the laboratory.

##### 1.2 Agentic AI Runtime (Edge/Cloud)

The LabOS vision-language model (VLM) ingests live streaming data combined with grounding inputs including reference standard operating procedures (SOPs), parameters measured by sensors or extracted visually from instruments. The system performs real-time step alignment, object and action tracking, error detection and correction, and generates time-stamped feedback for laboratory operations.

##### 1.3 XR Collaboration and Recording

System responses comprising visual overlays, voice guidance, and structured records are transmitted to XR devices in real-time. XR devices could render these visual and/or audio cues via an application (e.g. Unity or Android apps have been developed in this work). All decisions and supporting evidence are logged to tamper-evident run records ensuring reproducibility and audit compliance.

#### 2. On-Device XR Client Implementation

##### 2.1 Android and Unity Integration

The system integrates StreamKit (Android streaming framework) with a custom Unity application optimized for laboratory workflows. The Unity layer provides:

- An instruction player featuring step lists, timers, and parameter widgets
- A 3D annotation renderer for overlays anchored in 6-DoF space
- Raycast-to-surface capabilities for world-locked labels and bounding-box highlights

The client captures H.264/AAC or HEVC encoded video and audio, pose telemetry (position and orientation), and interaction events (voice commands, gaze focus, hand/gesture events). All streams are packetized with frame-accurate UTC timestamps and sequence identifiers.

##### 2.2 Bidirectional Command Channel

A persistent gRPC channel facilitates bidirectional communication:

- **Downlink**: Agent guidance including text, text-to-speech (TTS), overlay primitives, and corrective actions
- **Uplink**: Client status including SOP state, step progress, and device health metrics

Network endpoints (IP address and port) are configured through a lightweight JSON configuration file containing XR_AI_Runtime_IP and XR_AI_Runtime_PORT parameters.

### 3. Streaming and Edge Services Architecture

#### 3.1 Transport Pipeline Implementation

The on-device client publishes to an edge stream runtime supporting:

- **RTSP**: For video fan-out and network video recorder (NVR) integration
- **gRPC**: For low-latency bidirectional metadata exchange

Deployment includes stream pusher services, media servers, and optional NVR components orchestrated via Docker Compose for multi-camera, multi-user sessions. The pipeline maintains resilience to transient network interruptions through ring buffers on-device and back-pressure management at the edge, with recovered frames tagged as "late" but retained for audit purposes.

#### 3.2 Cloud Deployment Configuration

For cloud deployments, the LabOS agents can be hosted on various GPU inference platforms. XR clients transmit streams over WiFi to public endpoints with bidirectional guidance returned through duplex channels. The architecture maintains cloud provider independence through standardized interfaces.

### 4. Agentic AI Runtime Implementation

#### 4.1 Multimodal Input Processing

The system processes four categories of inputs:

1. **Live Streaming Data**: Video, audio, and pose information from XR and optionally, data from fixed cameras, equipment, sensors.
2. **Reference SOPs**: JSON-formatted protocols, SOPs, and key parameters with directed next pointers for step transitions.
3. **Grounding Signals**: Sensor data and visual extraction from devices and camera feeds.
4. **Configuration Parameters**: Routing information, session identifiers, and device addresses

#### 4.2 VLM-Driven Agentic AI Reasoning and Correction

LabOS VLM-powered agent processes structured clips with context to generate:

- Step identification (the step within protocol being performed)
- Status determination (correct/incorrect operation)
- Error type classification with affected parameters
- Minimal correction instructions
- Next step proposals upon completion

Responses undergo validation and prompt engineering to ensure structured responses are provided as feedback for the XR device / user log.

**5. Feedback Generation and XR Collaboration**

**5.1 Output Modality Specification**

- **Visual Overlays (3D)**: object labels and snap-to targets
- **Voice Guidance**: TTS-generated corrections and next actions
- **Temporal Tracking**: Live timelines annotating steps with a reference protocol, with tracker and time-point data for aligning on key parameters (such as temperature, timing/duration of the step).

**5.2 AI Assistance Features**

For high-precision operations enabled by Agentic AI, in addition to the Unity/Android app interface, the system can also project:

- Alignment guides derived from motion vectors.
- Indicators tied to SOP references for objects, actions, etc.

There are also hands-free control via voice, gaze, or gesture that enables step acknowledgment and repetition, reversal, adjustment requests.

**5.3 Bidirectional Streaming Implementation**

Feedback payloads (JSON with binary overlays) maintain budgeted latency targets. Unity clients render overlays in user frames with voice pre-buffered.

**6. Multi-User and Multi-Site Operation Capabilities**

The edge runtime supports N (currently up to 8) concurrent XR clients in team mode. Each stream receives a unique session and user identifiers with independent SOP state. A coordinator service could manage states for shared tasks. Cloud deployments enable sessions from distinct geographic sites to connect to common agent pools.

**7. Configuration and Session Initialization**

**7.1 Runtime Configuration**

XR clients load runtime IP addresses and ports from configuration files, establishing RTSP/gRPC/WebSocket connections to edge or cloud agents.

**7.2 SOP Loading**

JSON-formatted SOPs define hierarchical instruction nodes with documentation and key parameters. Directed progression is specified through next pointers.

**8. Example JSON file enabling laboratory protocol and parameters for LabOS XR-AI module alignment**

{

"title": "CRISPR CAS9 Delivery",
"id": "CAS9",
"version": "1.0",
"description": "Procedure for CRISPR CAS9 delivery",
"lab": {
"name": "Genetic Engineering Laboratory",
"department": "Molecular Biology",
"incharge_user": {
"name": "Le Cong Example",
"phone": "+1(617)863-6789",
},
"location": {
"address": "265 Campus Dr",
"city": "Stanford",
"state": "CA",
"country": "United States",
"zipcode": "94305"
}
},
"flowchart": {
"start_node": "1",
"nodes": [
{
"key": "1",
"value": {
"instruction": "Add 1 mL Opti-MEM into a sterile 1.5 mL EP tube.",
"description": "",
"type": "PROCESS",
"meta_data": {
"tts_audio_url": "",
"docs_url": "",
"video_url": "",
"picture_url": "",
"key_parameters": "temperature=37C, humidity=50, air_pressure=1013.25
hPa""""
},
"next": {
"default": "2"
},
"state": "NOT_STARTED"
}
},
{
"key": "2",
"value": {
"instruction": "Add Cas9 plasmid.",
"description": "",
"type": "PROCESS",

"meta_data": {
"tts_audio_url": "",
"docs_url": "https://labmanual.example.com/section1.1",
"video_url": "",
"picture_url": "",
"key_parameters": "temperature=37C, humidity=50, air_pressure=1013.25
hPa"""
},
"next": {
"default": "3"
},
"state": "NOT_STARTED"
}
},
{
"key": "3",
"value": {
"instruction": "Add guide RNA plasmid.",
"description": "",
"type": "PROCESS",
"meta_data": {
"tts_audio_url": "",
"docs_url": "",
"video_url": "",
"picture_url": "",
"key_parameters": "temperature=37C, humidity=50, air_pressure=1013.25
hPa"""
},
"next": {
"default": "4"
},
"state": "NOT_STARTED"
}
},
{
"key": "4",
"value": {
"instruction": "Add PEI (4:1 ratio to DNA).",
"description": "",
"type": "PROCESS",
"meta_data": {
"tts_audio_url": "",
"docs_url": "",
"video_url": "",
"picture_url": "",
"key_parameters": "temperature=37C, humidity=50, air_pressure=1013.25
hPa"""
},
"next": {

"default": "5"
},
"state": "NOT_STARTED"
}
},
{
"key": "5",
"value": {
"instruction": "Incubate at room temperature for 20 min.",
"description": "",
"type": "PROCESS",
"meta_data": {
"tts_audio_url": "",
"docs_url": "",
"video_url": "",
"picture_url": "",
"key_parameters": "temperature=37C, humidity=50, air_pressure=1013.25
hPa"""
},
"next": {
"default": "6"
},
"state": "NOT_STARTED"
}
},
{
"key": "6",
"value": {
"instruction": "Add the mixture dropwise into a 10 cm dish of 293T cells (~70-80%
confluency).",
"description": "",
"type": "PROCESS",
"meta_data": {
"tts_audio_url": "",
"docs_url": "",
"video_url": "",
"picture_url": "",
"key_parameters": "temperature=37C, humidity=50, air_pressure=1013.25
hPa"""
},
"next": {
"default": "7"
},
"state": "NOT_STARTED"
}
},
{
"key": "7",
"value": {

"instruction": "Gently rock the dish forward and backward to mix.",
"description": "",
"type": "PROCESS",
"meta_data": {
"tts_audio_url": "",
"docs_url": "",
"video_url": "",
"picture_url": "",
"key_parameters": "temperature=37C, humidity=50, air_pressure=1013.25 hPa"""
},
"next": {
"default": "8"
},
"state": "NOT_STARTED"
}
},
{
"key": "8",
"value": {
"instruction": "Place the dish back into the cell incubator (37°C, 5% $CO_2$).",
"description": "",
"type": "PROCESS",
"meta_data": {
"tts_audio_url": "",
"docs_url": "",
"video_url": "",
"picture_url": "",
"key_parameters": "temperature=37C, humidity=50, air_pressure=1013.25 hPa"""
},
"next": {
"default": "-1"
},
"state": "NOT_STARTED"
}
}
]
}
}
